# DIAFree enables untargeted open-search identification for Data-Independent Acquisition data

**DOI:** 10.1101/2020.08.30.274209

**Authors:** Iris Xu

**Affiliations:** Key Lab of Intelligent Information Processing of Chinese Academy of Sciences (CAS), Institute of Computing Technology, Chinese Academy of Sciences, Beijing 100190, China; University of Chinese Academy of Sciences, Beijing 100049, China

**Author notes:** To whom correspondence authors should be addressed, please contact.

## Abstract

As a reliable and high-throughput proteomics strategy, data-independent acquisition (DIA) has shown great potential for protein analysis. However, DIA also imposes stress on the data processing algorithm by generating complex multiplexed spectra. Traditionally, DIA data is processed using spectral libraries refined from experiment histories, which requires stable experiment conditions and additional runs. Furthermore, scientists still need to use library-free tools to generate spectral libraries from additional runs. To lessen those burdens, here we present DIAFree(https://github.com/xuesu/DIAFree), a library-free, tag-index-based software suite that enables both restrict search and open search on DIA data using the information of MS1 scans in a precursor-centric and spectrum-centric style. We validate the quality of detection by publicly available data. We further evaluate the quality of spectral libraries produced by DIAFree.

## Introduction

Proteins are the cornerstones and regulators of lives. A powerful strategy to identify and quantify protein in large-scale experiments is liquid chromatography tandem mass spectrometry (LC-MS/MS). Currently, the most popular approach of LC-MS/MS remains data-dependent analysis (DDA). However, in the DDA method, the spectrometer needs to choose precursor ions that should be isolated and fragmented, which, in consequence, ignores some signals of low-abundance peptides, induces uncertainty and impairs the quantitative stability of tandem spectra (MS2). To alleviate this problem, scientists proposed the data-independent acquisition (DIA) method(1, 2). Instead of investigating one precursor ion in one MS2 spectrum, in the DIA method, the spectrometer fragments all precursor ions within a user-specified mass-to-charge ratio(m/z) isolation window.

However, the DIA method also imposes challenges on protein identification algorithms by generating complex multiplexed spectra. Popular tools, such as OpenSWATH(3), Skyline(4-6), DIA-NN(7) and EncyclopeDIA(8), use the spectral libraries built from historical data(9), which often contain the information of fragmentation patterns, retention time, and chromatography patterns(8), to extract peptide signals from complex spectra. However, despite higher sensitiveness and faster speed, the library-based algorithm has three lapses(9-11). First, the algorithm can only identify analytes inside the library. Second, additional runs or strictly invariant experiment conditions are required to keep the library effective. Third, iterative library searches may accumulate errors in the spectral library.

In some studies, machine learning methods, such as pDeep(12, 13) and Prosit(14, 15), is employed to lessen those limitations and generate spectral libraries(7, 14, 16). Still, the presence of post-translational modification (PTM) generates many peptidoforms(17-19), which makes the predicting process time-consuming. Therefore, we still need complementary library-free methods during peptide identification.

A group of library-free software such as DIA-Umpire(11) and Group-DIA(20) demultiplex coeluting spectra to generate multiple pseudo MS2 spectra that contain fewer ion interferences utilizing information such as MS1 scans, extracted ion chromatograms (XICs) of precursor ions and isolation window settings. After that, software that developed for DDA data can directly analyze such pseudo spectra. The demultiplexed method can improve the signal-to-noise ratio, expose low-abundance peptide, and accelerate the speed of identification. However, this method might hide fragment ions that are affected by coelution, which leads to lower sensitiveness in low-quality DIA data. Other library-free algorithms employ DIA-specific properties to assess the existing evidence of peptides. For example, PEAKS(21) directly inputs the adjacent spectra into a neural network. PECAN(22) uses the fragment extracted ion chromatograms (XICs) to qualifying the evidence of peptides. However, those algorithms do not guarantee the precursor evidence of identified peptide. Since that too many fragment ions signals do not belong to the same peptide but have similar XICs, orphan peptides(23) identified by those signals are questionable. This problem is especially significant in the open search mode.

In this article, we proposed an untargeted library-free open-source software suite DIAFree. The DIAFree workflow qualifies peptides by assessing the cohesion of fragment ions, the similarity of XICs between the precursor and fragments, and the quality of the match between the peptide and the spectrum, which eliminates the need for spectral library or retention time information. The DIAFree workflow enables ordinary restrict search along with open search, which means that DIAFree can search more than 1,000 kinds of unexpected modification as well as user-defined modification. We believe that the open search can help scientists explore potential modifications. In the DIAFree workflow, we first extract precursors and the XICs of precursors using theoretical isotope pattern, then use the tag index generated by Open-pFind(24) to discover peptide candidates. After that, we use a feature-grouped, semi-supervised learning algorithm which composed of XG-Boost sub-classifiers to rerank those candidates. The DIAFree workflow employs both spectrum-centric and precursor-centric strategies to secure accuracy, which means the workflow only selects one peptide candidate from each precursor and discards highly similar peptides in each spectrum. Last, we provide an interactive result viewer that enables manual validation and refinement.

## Methods

### The DIAFree Workflow

The workflow has several components, as shown in Figure 1. First, the pXtractDIA converts raw MS/MS data (.mzML or .raw) into an SQLite database. Then the DIAFreeParse reconstructs potential isotope clusters from MS1scans and then extracts putative precursor ions. Instead of considering unfragmented precursor ion signals in MS2, we only accept guaranteed isotope clusters from MS1 scans to avoid false-positive peptides, since too many peaks in complex MS2 scans are not related but have similar XICs. The quality of the isotope cluster is assessed by comparing this reconstructed isotope cluster to the theoretical cluster calculated by the Averagine(25) model using a simplified EMASS(26) algorithm implementation. In this article, we call a virtual isotope as a non-existing isotope peak that has smaller neuron numbers than real mono isotope. Virtual isotopes have high quality when an isotope is misidentified as mono; therefore, we also restrict the quality of virtual isotopes in identified isotope clusters to avoid precursors that are from isotope peaks of others.

**Figure 1.**
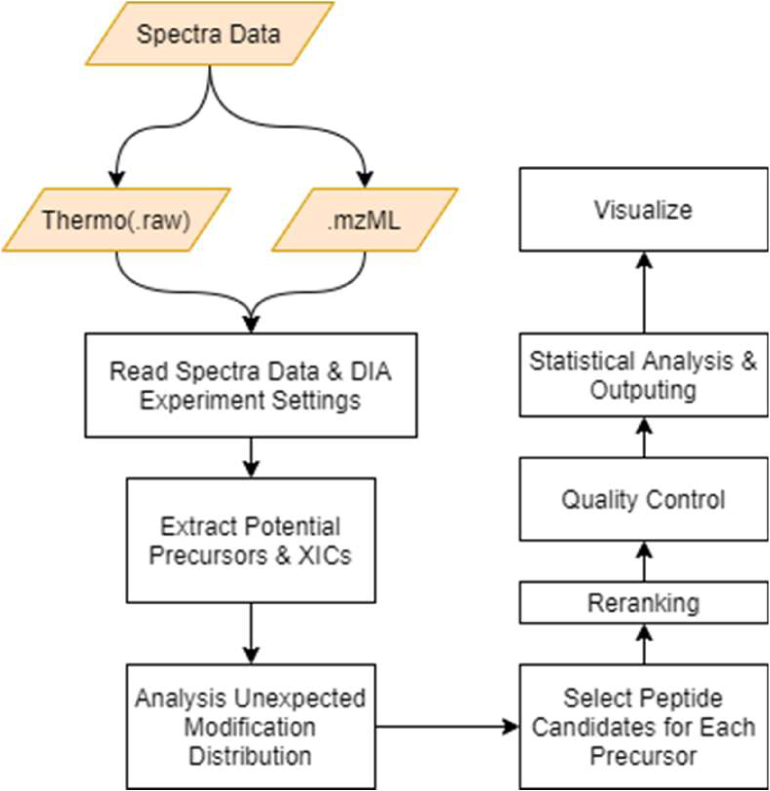
The workflow of DIAFree

For each precursor, the DIAFree then selects out all putative Peptide-Spectrum Match (PSM)s from the intermediate result of Open-pFind(24). The tag index used by Open-pFind ensures all putative PSMs have at least five continuous fragment ion signals matched in the open search mode, which can reduce the size of the search space. To speed up, we prohibit rating and reranking from Open-pFind. After that, DIAFreePostProcess generates features and the main score for each putative PSM and filters out less-reliable putative PSMs. After that, DIAFreePostProcess employs a feature-grouped, semi-supervised learning algorithm derived from Percolator(27) to rerank those putative PSMs. The learning algorithm has two key innovations to avoid overfitting. First, rather than using one support vector machine (SVM), we ensemble three different XGBoost(28) sub classifiers. The reason for using XGBoost instead of SVM is because XGBoost has the superior ability to handle unbalanced samples. Each sub classifier is trained with different features and aims: the first classifier is from the view of how good a putative peptide matches the MS2 scan at the apex of the XIC, the second classifier to evaluate the similarity of XICs between fragment ions and the precursor ion, the third classifier to assess the cohesion of fragment ions. A sub classifier using theoretical ion distributions generated by pDeep, if specified by the user, is also supported. Second, in the first round of training, positive and negative samples are simply determined by the main scores. However, in the following rounds, only those samples that approved by more sub classifiers can be used. By using this training strategy, we can prevent learning from less-confident samples. We can also limit the results reported by DIAFree, which match an MS2 scan well but are weak in the chromatogram-related properties.

Manual analysis is crucial to help researchers discover new PTMs and new proteins. Here we implement DIAFreeViewer to visualize and validate the result. Aside from visualizing the peptides and the proteins, users can also retrieve high-quality peptides by score threshold that are ruled out by precursor-centric quality control or spectrum-centric quality control. The DIAFreeViewer enables manually refining the retention time boundary of the XIC and then re-calculating the relative abundance of peptides. The viewer also supports exporting selected peptides into a spectral library.

### Read spectra data and experiment settings

The DIAFreeParse extracts spectra and experiment settings from the data (.raw or .mzML). Profile peaks, if provided, are centralized while reading. For AB SCIEX data, the pXtractDIA further denoises those spectra by discard peaks whose signal intensities are below a certain threshold (0.01 * strongest intensity in the same scan for MS1, 0.03 for MS2) and using a dynamic planning algorithm inspired by BackgroundDetector algorithm from DIA-Umpire(11) with the ratio between the number of isotopes and the number of noises greater than 1.

In this paper, data collected from AB SCIEX instruments were transformed into mzML using the ProteoWizard package (version: 3.0.20034) with the following settings:

--filter “peakPicking vendor msLevel=1-2” --filter “threshold absolute 2 most-intense”

Dataset ENC_yeast_board were transformed into mzML with the following settings:

--filter “peakPicking vendor msLevel=1-2” –demultiplex “optimization=overlap_only massError=10.0ppm”

### Extract potential precursors and XICs

For each putative isotope cluster, features of the isotope pattern and XICs are reconstructed by DIAFreeParse and sent to a pre-trained MARSpline(29) model to examine if the cluster is a qualified isotope cluster. Precursors are then extracted and sent to Open-pFind with the information about the MS2 scan at the apex of the mono XIC. The DIAFreeParse also outputs chromatograms for the following procedures.

Here we request that all putative reconstructed isotope clusters sent to MARSpline must have mono isotopes and 1st isotopes presented. The similarities of XICs between any virtual isotope and real isotopes must be lower than the similarities between the mono isotope and other real isotopes. In DIAFreeParse, the Averagine(25) model obtained from the PIR protein database (C_4.9384_H_7.7583_N_1.3577_O_1.4773_S_0.0417_) was chosen to generate a theoretical formula for each possible ion mass(0-12000Da). Users can also customize this Averagine amino acid formula. A simplified EMASS(26) algorithm based on dynamic planning was then used to calculate the isotope pattern of the formula. We only accept reconstructed isotope clusters with a correlation coefficient higher than a certain threshold (here, 0.6 was used) between theoretical isotope pattern and reconstructed pattern. We use two virtual isotopes and five real isotopes in the model. The DIAFreeParse uses the Hash-induced Linked List to accelerate the chromatogram extraction. The MARSpline model was trained by three DDA datasets: Mann-Hela(30), Gygi-Human(31) and Mann-Mouse(32). Here we avoid using DIA datasets to prevent ion inference accumulation.

### Selecting putative peptides

After removing the redundant PSMs, data are sent to Open-pFind to select out putative peptides. Here we use Open-pFind to free from the labor of enumerating protein substrings in DIAFreePostProcess. Reranking PSMs^1^ and Reducing protein database in Open-pFind are disabled. We keep the top 30 PSMs for each precursor from the intermediate results of Open-pFind.

### Preprocessing Spectrum

For each MS2, DIAFreePostProcess removes all isotope peaks in a reliable isotope cluster whose correlation coefficient is above a threshold (in this paper, 0.8 was used) between this cluster and the theoretical isotope pattern. After deconvolution, for each precursor, the peaks corresponding to the precursor ion, its isotopes and its neutral losses are also removed in MS2.

### Quality Control

The DIAFreePostProcess applied several restrictions on the PSMs to reduce the number of false-positive peptides reported.

1. The peptide must have at least six fragment ions matched in the original MS2 scan at the apex.
2. The peptide must have at least four fragment ions whose PCC is above 0.7 when compared with the XIC of the median peak trace and have at least two fragment ions whose PCC is greater than 0.7 when compared with the XIC of the precursor.
3. PCC between the XIC of the precursor and the median peak trace must be greater than or equal to 0.5.
4. The XIC of the precursor, the XIC of the median peak trace and the overlap between those XICs must be longer than one cycle.

A restriction is specified to control unexcepted modifications.

1. Let S0_p_ = S_p_ + log(UMod!), where S_p_ is the main score of this PSM. We require that the ratio of S0_p_ contributed by fragment ions must greater than or equal to 0.1.
2. PCC between the median peak trace of fragment ions that are affected by unexpected modification and fragment ions that are unaffected must be greater than or equal to 0.7.

Users can customize the restrictions in DIAFree. We select out ten strongest fragment ions (except y1+, b1+, y1++, b1++) and then calculate the median peak trace in a similar way of EncyclopeDIA(8) used, except that we use linear interpolation. We use the same method to calculate the median peak trace with affected fragment ions and trace with unaffected fragment ions.

### Main Score of DIAFreePostProcess

The main score of DIAFreePostProcess is based on Two-Poisson Model(33). As a scoring method, this main score is easy to understand and compute. For a peptide p, the main score *S*_p_ is calculated as:

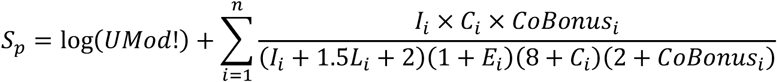

where *UMod* is the number of unexpected modifications in the peptide, *n* is the number of fragment ions, *I*_*i*_ is the normalized and triangular-smoothed intensity of i_th_ fragment ion, *C*_*i*_ is the correlation coefficient between the XIC of precursor and the fragment ion, *E*_*i*_ is the normalized fragment error, *L*_*i*_ is the pivot-normed peptide length. If we name the triangular-smoothed MS2 scan at the apex of the XIC as *ms*2, then *I*_*i*_ is calculated as:

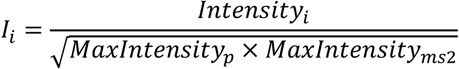

Here the continuity bonus *CoBonus*_*i*_ is set as:

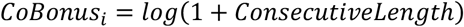

Where the consecutive length is the number of matched consecutive fragment pairs whose PCC compared with the XIC of the precursor are greater than 0.7.

### Reranking

A semi-supervised learning method is used to iteratively separate target PSMs from decoy ones based on XGBoost. To avoid overfitting, three different groups of features are distributed to three different sub classifiers to keep all sub classifiers know only part of features and must rely on the know information:

1. ShortTerm: Features generated from 1-3 MS2 scans near the apex of precursor chromatogram. The ShortTerm sub classifier is focusing on the matching degree between the peptide and those MS2 scans.
2. LongTerm: Features related to the XIC of the precursor.
3. LongTermInner: Features related to the cohesion of the fragment ions. Most features are related to the median peak trace.

By default, the XGBoost classifiers are at most four layers depth with L2-regularization enabled. During training, the scalePosWeight is set as the ratio of the number of negative samples to the positive samples. The learning rate of XGBoost is 0.1.

If pDeep-related features are used, DIAFreePostProcess first extracts all putative peptides passed quality control, turns those peptides into the corresponding forms without modifications (except phosphorylation), removes duplicate peptides, then sends those new peptides to pDeep and stores the output into an SQLite database. DIA-based refinement is banned to avoid ion interference.

For each round, sub classifiers vote for the negative and positive training samples. Each sub classifier selects one best peptide match from one precursor as training sample candidates. For each sub classifier, target PSMs within the threshold of 5% PSM FDR and the threshold of 1% Peptide FDR are marked as positive training samples; target PSMs above the threshold of 20% FDR and decoy PSMs are marked as negative training samples. If a PSM is marked by at least half of sub classifiers as positive and has more positive votes than negative votes, this match is identified as a positive training sample. If all sub classifiers mark a PSM as negative, this match is then selected as a negative sample.

The final scores are obtained by simply averaging all sub scores (SimpleAvg).

### Chromatogram Refinement

The DIAFree workflow refines precursor chromatogram at two steps: First, when DIAFreePostProcess reads the chromatograms, the software re-extracts those chromatograms to avoid actual isotope m/z drifts away from the theoretical m/z. Second, before quantification, we refine the peak boundaries using the median peak trace.

### Protein Inference

Protein inference methods used by protein identification software are diverse. Here we implement a simple protein inference method that derived from the inference method in Open-pFind. Before calculating the FDR, proteins are divided into protein groups using supporting peptides. After that, we calculate several features for each protein group and FDR correct those protein groups with a reimplementation of Percolator(27). Those features include the coverage of protein, the sum of the best PSM final score from each unique peptide, from each peptide, and from each amino acid sequence.

### Statistical Analysis

As we described above, only one best peptide is kept for each precursor. If a pair of similar peptides present in the same apex MS2 scan (The overlap of sequence between this pair of peptides >= 90% * the length of shorter peptide), DIAFreePostProcess only keep the peptides with at least one independent ion evidence or higher final score. However, users can still retrieve, view, and output all those eliminated high-quality peptides in DIAFreeViewer.

We use native FDR since it is the most stable method. When the target library is too powerful to mix up with decoys on the 1% FDR level, methods like pyProphet(19, 34) are unable to estimate π_O_. Adding more decoys or provide high quality decoy entries in spectral library should be a solution; however, more decoys may violate the internal assumption of statistical analysis software. Still, users can output Percolator(27)-compatible results (.tsv) along with TPP(35, 36)-compatible results (.pepXML) using DIAFree.

### Quantification

In DIAFree, we provide a quantification schema, including several ordered layers: transitions, PSMs, peptides, peptides with unlocalized modifications, peptide sequences, and proteins. We also offer a series of user-defined settings, including 1. The methods of estimating the abundance of an element in each layer. 2. The number of elements needed to quantify an element in the next layer. 3. The methods of ranking elements, which can determine which element should be further passed to the next layer with the limitation of the number needed. 4. The methods of judging whether an element should be prohibited in the quantification of the next layer. 5. The whitening methods of each layer. Those settings enable us to implement different quantification style including DIAUmpire-Style and EncyclopeDIA-Style. For a labeled experiment, we also support calling pQuant(37) (MS1) or pIsobaric (MS2) to infer the abundance.

### Manual Validation and Refinement

First, in DIAFreeViewer, users can visualize score distribution, spectral and chromatographic data plots of results. Users can also modify the chromatography boundary of a precursor and recalculate the score of all putative PSMs with the help of models generated by DIAFreePostProcess. Besides, if users input a peptide sequence or a protein sequence, all peptidoforms, with main scores and final scores above certain thresholds, are searched and displayed by DIAFreeViewer. Furthermore, DIAFreeViewer allows users to selectively output results. The DIAFreeViewer uses lazy loading and fixed-size caching to support visualization of large-scale experiments.

### Software Runtime

All the experiments were performed on a DELL G5 5090 with Window 10, Inter Core™ i7-9700 CPU (3.00GHz), 80.0GB RAM and a 5400 RPM WD hard disk. In order to prevent inferences, we run each experiment separately and pay extra attention to not run any other program during experiments.

### Hela SILAC Peptides Preparation

In normal conditions, Hela cell was cultured in Dulbecco’s Modified Eagle Medium (DMEM) medium supplemented with 10% fetal bovine serum (FBS). For SILAC experiment, the heavy isotopically labeled Hela cells were prepared according to the previous described protocol(38). The heavy SILAC medium was prepared by adding heavy isotopically amino acids, 13C615N2 L-Lysine-2HCl (ThermoFisher, 89988) and 13C615N4 L-Arginine-HCl (ThermoFisher, 89990), and 10% dialyzed fetal bovine serum (ThermoFisher, 30067334) to DMEM deficient in both L-lysine and L-arginine for SILAC (ThermoFisher, A33822). The Hela cell was cultured for eight passages in heavy SILAC medium to reach a nearly complete labeling (>99%, checked by MS).

Light and heavy SILAC Hela cells were grown to confluence, and then collected by washing with precooled PBS thrice and snap-frozen in liquid nitrogen. The cells were lysed with SDS lysis buffer (4% SDS, 100 mM Tris-HCl, 0.1 M DTT, pH 7.6) and kept in −80°C for the following proteomic analysis. For protein extraction, the light and heavy cells were sonicated at 15% amplitude for 5 s on and 5 s off with the total working time of 1 min (JY92-IIDN, Ningblio Scientz Biotechnology Co., LTD, China). The proteins were then denatured and reduced at 95°C for 5 min. The insoluble debris was removed by centrifugation at 12,000 g for 10 min and the supernatant was retained for proteomic experiment. The protein concentration was determined using tryptophan-based fluorescence quantification method. Eventually, the light and heavy SILAC proteins were mixed in equal amounts.

Filter-aided sample preparation (FASP) procedure was used for protein digestion. Briefly, proteins were loaded in 10 kDa centrifugal filter tubes (Millipore), washed twice with 200 μL UA buffer (8 M urea in 0.1 M Tris-HCl, pH 8.5), alkylated with 50 mM iodoacetamide in 200 μL UA buffer for 30 min in the darkness, washed thrice with 100 μL UA buffer again and finally washed thrice with 100 μL 100 mM NH4HCO3. All above steps were centrifuged at 12,000 g at 25°C. Proteins were digested at 37°C for 18 hr with trypsin (Promega) at a concentration of 1:25 (w/w) in 100 mM NH4HCO3. After digestion, peptides were eluted by centrifugation. The pH of the peptides solution was adjusted to < 2.0 by adding pure formic acid and subjected to C18 solid-phase extraction desalting (Waters, Sep-Pak® Vac 3cc 200 mg tC18 Cartridge). The amount of the purified peptides was determined using Nanodrop One (Thermo Scientific).

### DIA MS Data Acquisition

The Hela SILAC peptides were resolved using 0.1% formic acid and 1 μg peptides were injected per MS run. The peptides were separated using a home-made micro-tip C18 column (75 mm × 200 mm) packed with ReproSil-Pur C18-AQ, 1.9 μm resin (Dr. Maisch GmbH, Germany) on a nanoflow HPLC Easy-nLC 1200 system (Thermo Fisher Scientific), using a 120 min LC gradient at 300 nL/min. Buffer A consisted of 0.1% (v/v) formic acid in H2O and Buffer B consisted of 0.1% (v/v) formic acid in 80% acetonitrile. The gradient was set as follows: 2%–5% B in 1 min; 5%–33% B in 93 min; 33%– 45% B in 15 min; 45%–100% B in 3 min; 100% B in 8 min.

Proteomic analyses were performed on a Q Exactive HF mass spectrometer (Thermo Fisher Scientific). The spray voltage was set at 2,300 V in positive ion mode and the ion transfer tube temperature was set at 300°C. Data-independent acquisition was performed using Xcalibur software in profile spectrum data type. The MS1 full scan was set at a resolution of 120,000 @ m/z 200, AGC target 3e6 and maximum IT 100 ms by orbitrap mass analyzer (350-1650 m/z), followed by DIA MS2 scans generated by HCD fragmentation at a resolution of 30,000 @ m/z 200, AGC target 5e5 and auto maximum IT. The fixed first mass of MS2 spectrum was set 100.0 m/z. The normalized collision energy (NCE) was set at NCE 27%. The RT windowed DIA method was introduced previously(39). The DIA mass window settings for SILAC_20Da, SILAC_5Da and SILAC_RTWin were described as follows: 1) SILAC_20Da, 350-1010 m/z, window size 20 Da, 33 windows; 2) SILAC_5Da, 4 gas phase fractionated fractions, 350-500 m/z, window size 5 Da, 30 windows; 500-650 m/z, window size 5 Da, 30 windows; 650-800 m/z, window size 5 Da, 30 windows; 800-1000 m/z, window size 5 Da, 40 windows; 3) SILAC_RTWin, 400-600 m/z in 0-60 min; 600-800 m/z in 60-90 min; 800-1000 m/z in 90-120 min, window size 5 Da, 40 windows. Three technical replicates were analyzed for each condition.

## Results

### Restrict Search using DIAFree

We assessed the result of DIAFree by benchmarking DIAFree against other widely used sequence database search tools including Walnut(22) (a reimplementation of PECAN, carried by EncyclopeDIA-0.9.0) and a pipeline of DIAUmpire(11)(Release at 2016-04-24), Comet(40) (41)(version: 2019013)and Percolator(27)(version: 3.0.1). Datasets used (details are shown in Supplementary Table 1) are ranging from lower complexity (UPS1 standard and UPS2 standard, each contains 48 human proteins) to higher complexity (E.coli, yeast, and human). We have observed that DIAFree can report more peptide sequences (4.39% - 84.56% more peptide sequences compared to Walnut, 4.06%-99.46% more peptide sequences compared to DIAUmpire) with less running time (Walnut used 39%-216% more time than DIAFree, DIAUmpire used 0.7%-1656% more time than DIAFree). Since a peptide sequence identified by both the DIA dataset and the corresponding DDA dataset is more reliable than those cannot, we also searched for DDA data using Open-pFind for each sample. We combined multiple DDA datasets that come from the same biological sample into one large dataset. The DDA coverage rate of a DIA dataset is defined as the number of identified peptide sequences from this dataset that can also be identified by using the corresponding DDA dataset divided the total number of identified sequences in this DIA dataset. We found that most peptide sequences identified by DIAFree using DIA data were also identified by the corresponding DDA data (DDA coverage rate is between 83.39% to 92.50% in the dataset where the number of identified sequences in the corresponding DDA dataset is larger than the number in this DIA dataset). We then used an entrapment strategy to evaluate the quality of detected peptides. We mixed Swiss_human sequence library with Swiss_yeast sequence library and use the nearly 1: 1 mixed library to search Hela (SILAC_Hela_RTwin, SILAC_Hela_20Da, SILAC_Hela_5Da, PECAN_1xGPF, PECAN_2xGPF, PECAN_4xGPF) and yeast data (ENC_yeast_board, ENC_yeast_narrow). We found that DIAFree has smaller ratios of entrapment peptides in most datasets. Datasets, including SILAC_Hela_RTwin, SILAC_Hela_20Da, and SILAC_Hela_5Da, have been labeled by SILAC method and were also used to assess the identification precision. We believe that given an identified light-labeled peptide, a corresponding SILAC heavy-labeled peptide ion likely exists in the same scan. Therefore, if we cannot find the precursor evidence and the fragment evidence of a corresponding heavy-labeled peptide, this result is unreliable. Here the precursor evidence of a peptide is regarded as existing when both a light-labeled and a heavy-labeled precursor ion in the same charge state can be matched by the leading MS1 scan or the following MS1 of the apex MS2 scan. The fragment evidence of a peptide is defined in the same way: If the heavy-labeled fragment ion and the light-labeled fragment ion exist in the apex MS2 scan, and this pair of fragment ions have been affected by SILAC and have different m/z, we call this pair of fragment ions as fragment evidence of a peptide. After selecting one best scoring PSM for each peptide sequences, we found that at least 99.44% identified sequences in all datasets have precursor evidence and at least one pairs of fragment ion evidence. Furthermore, we use a popular spectral library search tool EncyclopeDIA(8) to search dataset PECAN_1xGPF and SILAC_Hela_20Da against a Hela-based spectral library panhuman.dlib(8). After that, we draw a scatter plot for each dataset using those identified precursor ions that are identified by both DIAFree and EncyclopeDIA (18808/20278 precursor ions that identified by DIAFree in the dataset PECAN_1xGPF, 18659/21885 in the SILAC_Hela_20Da). Although DIAFree do not use any information of peptide retention time in the sequence searching mode, we found that the retention time of identified PSMs is highly correlated with the retention time of the corrected retention time in the EncyclopeDIA ELIB library and is highly reliable. Last, we searched datasets with spectral libraries generated by DIAFree using EncyclopeDIA. We found that most precursor ions in the spectral libraries can be re-identified by EncyclopeDIA again (95.67% precursor ions are identified in the DIAUmpire_Ecoli, at least 99% precursor ions in other datasets).

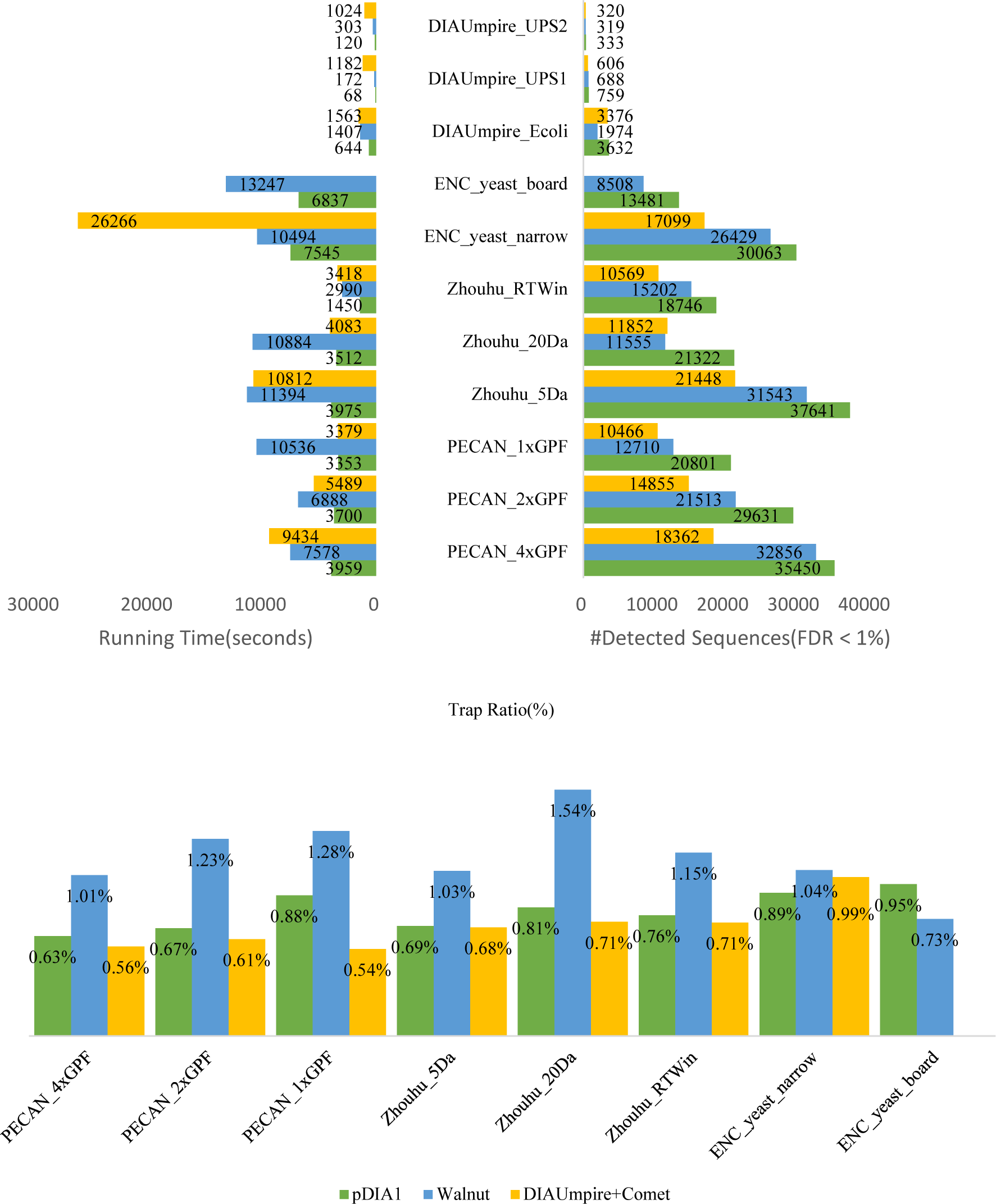

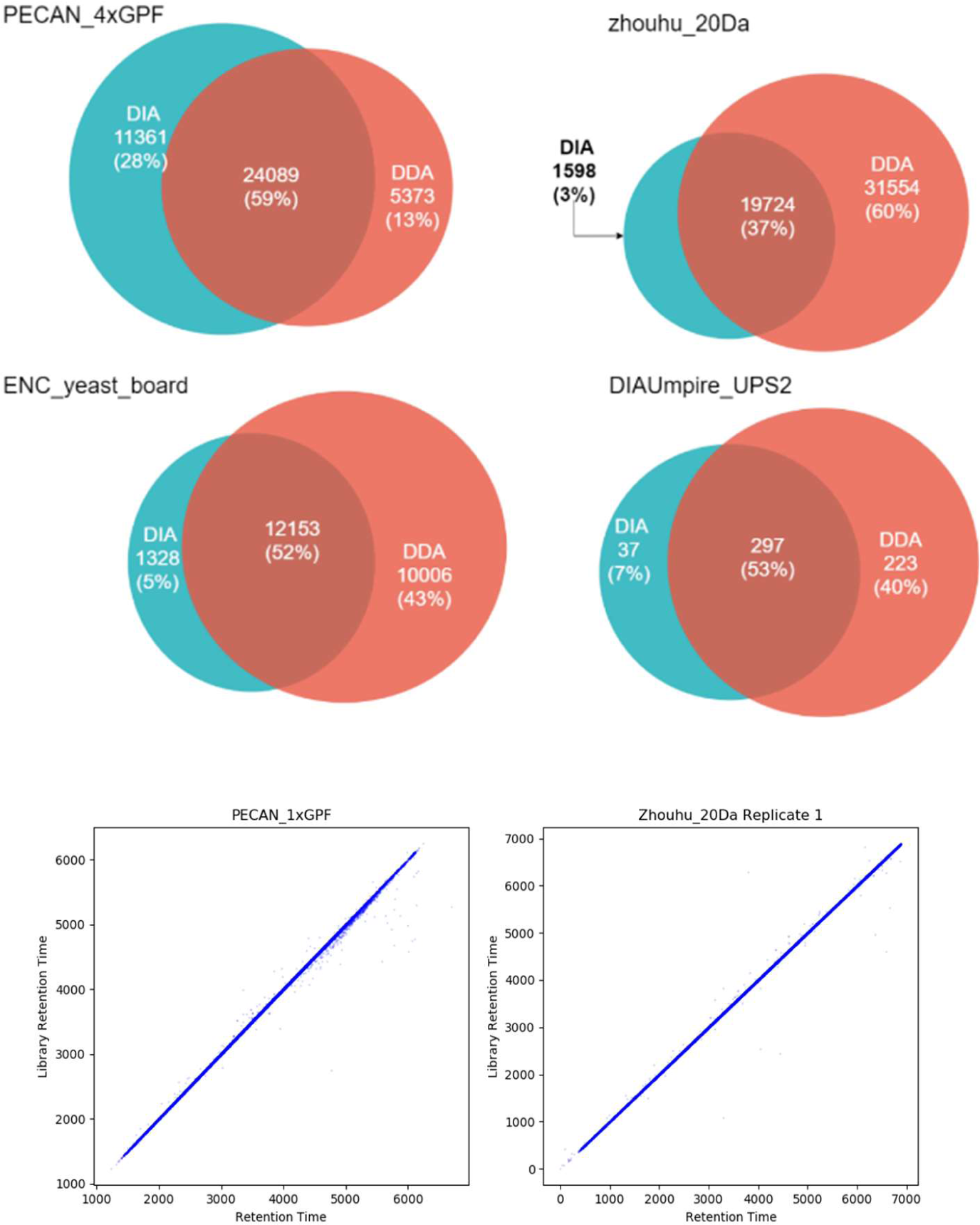

### Open Search using DIAFree

We define the unexpected modification as modifications that are neither set as a fixed modification nor variable modifications. Open Search is a feasible method for discovering frequent unexpected modifications or single amino acid mutations in spectra. Here we set Carbamidomethyl as a fixed modification, Oxidation as variable modification, disabled shrinking protein sequence library by first search pass that estimates the distribution of all modifications (salvo_iteration=0), and then searched each dataset using open search mode. We regard modifications that carried by ⩾ 50 PSMs as high-frequency modifications. We found that high-frequency modifications between each replicate in each dataset are similar. Here Glu->Phe was discovered in all three replicates, but in Replicate2, the PSM frequency of this modification is only 45; in Replicate3, the frequency is 47. Both frequencies are not far from 50.

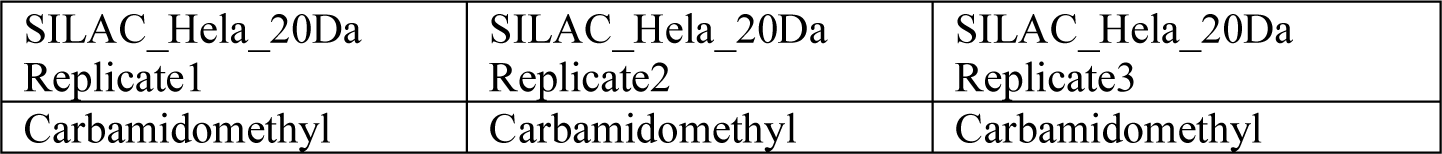
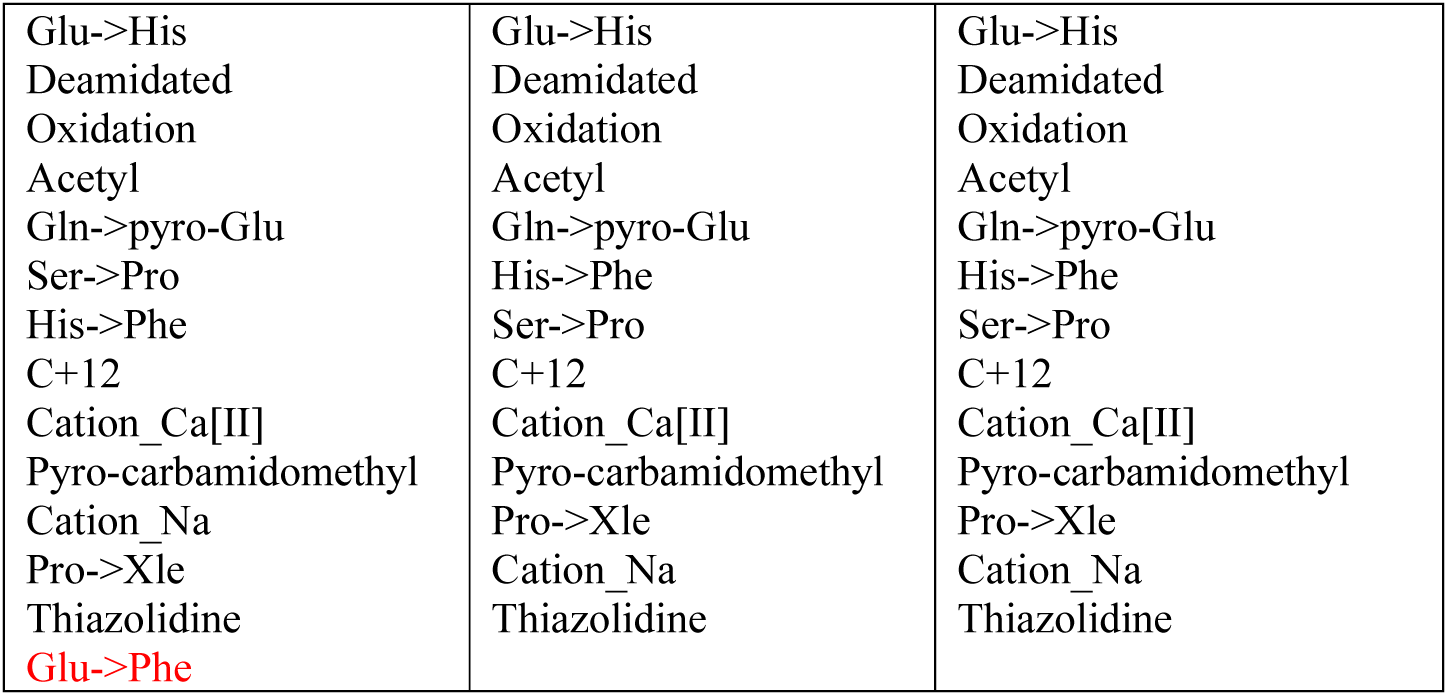

|  |  |  |
| --- | --- | --- |
| SILAC_Hela_20Da<br>Replicate1 | SILAC_Hela_20Da<br>Replicate2 | SILAC_Hela_20Da<br>Replicate3 |
| Carbamidomethyl | Carbamidomethyl | Carbamidomethyl |
| Glu->His<br>Deamidated<br>Oxidation<br>Acetyl<br>Gln->pyro-Glu<br>Ser->Pro<br>His->Phe<br>C+12<br>Cation_Ca[II]<br>Pyro-carbamidomethyl<br>Cation_Na<br>Pro->Xle<br>Thiazolidine<br>Glu->Phe | Glu->His<br>Deamidated<br>Oxidation<br>Acetyl<br>Gln->pyro-Glu<br>His->Phe<br>Ser->Pro<br>C+12<br>Cation_Ca[II]<br>Pyro-carbamidomethyl<br>Pro->Xle<br>Cation_Na<br>Thiazolidine | Glu->His<br>Deamidated<br>Oxidation<br>Acetyl<br>Gln->pyro-Glu<br>His->Phe<br>Ser->Pro<br>C+12<br>Cation_Ca[II]<br>Pyro-carbamidomethyl<br>Pro->Xle<br>Cation_Na<br>Thiazolidine |

Furthermore, high-frequency modifications in datasets with more fractions can cover modifications in datasets collected from the same sample but have fewer fractions.

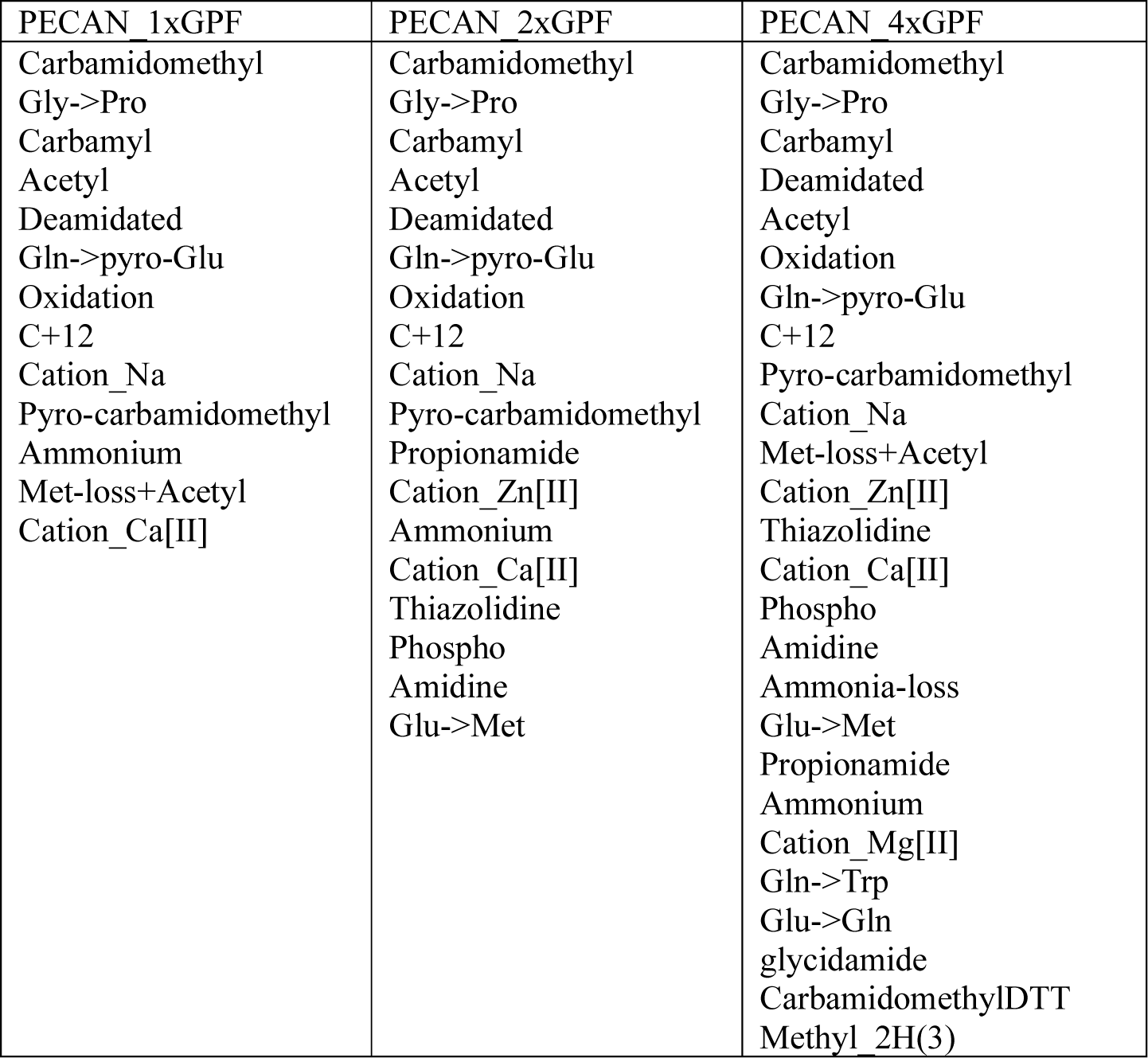

| PECAN 1xGPF | PECAN 2xGPF | PECAN 4xGPF |
| --- | --- | --- |
| Carbamidomethyl<br>Gly->Pro<br>Carbamyl<br>Acetyl<br>Deamidated<br>Gln->pyro-Glu<br>Oxidation<br>C+12<br>Cation_Na<br>Pyro-carbamidomethyl<br>Ammonium<br>Met-loss+Acetyl<br>Cation_Ca[II] | Carbamidomethyl<br>Gly->Pro<br>Carbamyl<br>Acetyl<br>Deamidated<br>Gln->pyro-Glu<br>Oxidation<br>C+12<br>Cation_Na<br>Pyro-carbamidomethyl<br>Propionamide<br>Cation_Zn[II]<br>Ammonium<br>Cation_Ca[II]<br>Thiazolidine<br>Phospho<br>Amidine<br>Glu->Met | Carbamidomethyl<br>Gly->Pro<br>Carbamyl<br>Deamidated<br>Acetyl<br>Oxidation<br>Gln->pyro-Glu<br>C+12<br>Pyro-carbamidomethyl<br>Cation_Na<br>Met-loss+Acetyl<br>Cation_Zn[II]<br>Thiazolidine<br>Cation_Ca[II]<br>Phospho<br>Amidine<br>Ammonia-loss<br>Glu->Met<br>Propionamide<br>Ammonium<br>Cation_Mg[II]<br>Gln->Trp<br>Glu->Gln<br>glycidamide<br>CarbamidomethylDTT<br>Methyl 2H(3) |

Using those datasets, we summarized top-10 high-frequency modifications that occurred in each sample. We discovered the evidence of phosphorylation and glycosylation from UPS1/UPS2 datasets as expecting yet with low frequency.

| HeLa | yeast | E. coli | UPS1 |
| --- | --- | --- | --- |
| Pyro-carbamidomethyl<br>Oxidation<br>Gln->pyro-Glu<br>Deamidated | Carbamidomethyl<br>Carbamyl<br>Oxidation<br>Deamidated | Carbamidomethyl<br>Deamidated<br>Oxidation<br>Gln->pyro-Glu | Oxidation<br>Deamidated<br>Carbamidomethyl<br>Gln->pyro-Glu |
| Carbamidomethyl<br>C+12<br>Acetyl<br>Single amino acid<br>mutation (Glu->His,<br>His->Phe, Ser->Pro)<br>Cation_Na<br>Cation_Ca[II] | Acetyl<br>Gln->pyro-Glu<br>Cation_Na<br>Met-loss+Acetyl<br>Cation_Ca[II]<br>Ammonia-loss<br>Propionamide |  |  |

With open search, more peptides can be identified. To validate the quality of those results, we also used the entrapment strategy in this section. As the figure shows, the ratio of entrapment peptides in all datasets are still below 2% but higher than the ratios in the restricted search, which is because that peptides with unexpected modifications have a higher chance to connect originally unrelated two groups of fragment ions into one PSM by a low-frequency unexpected modification and therefore have a higher entrapment ratio. We also applied SILAC validation on open search results by selecting the best PSM for each peptide. In all SILAC-labeled datasets (SILAC_Hela_20Da, SILAC_Hela_5Da, SILAC_Hela_RTWin), at least 94.25% peptide that affected by SILAC label have precursor evidence and at least a pair of fragment evidence.

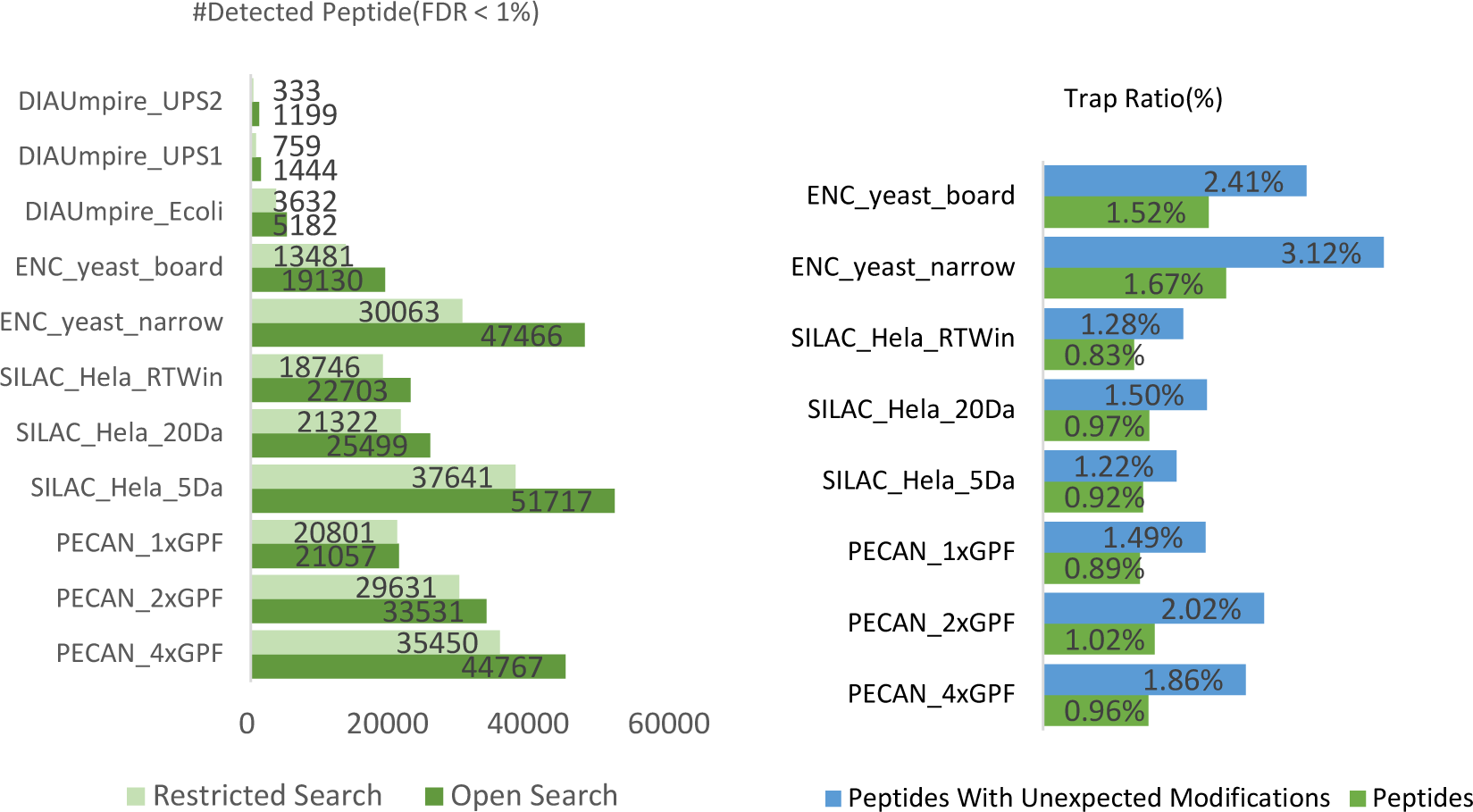

## Support information

**Supplementary Table 1.**
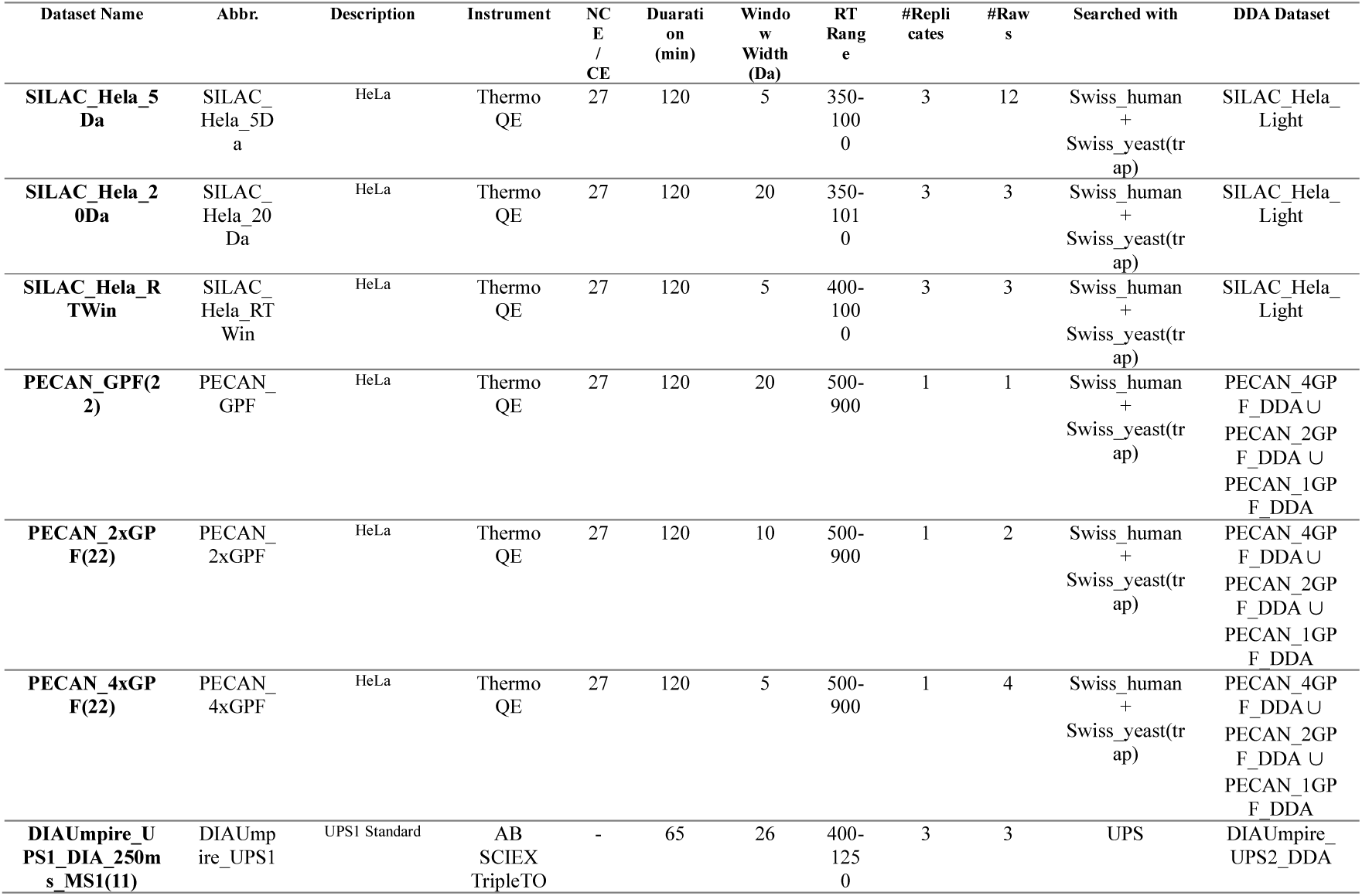
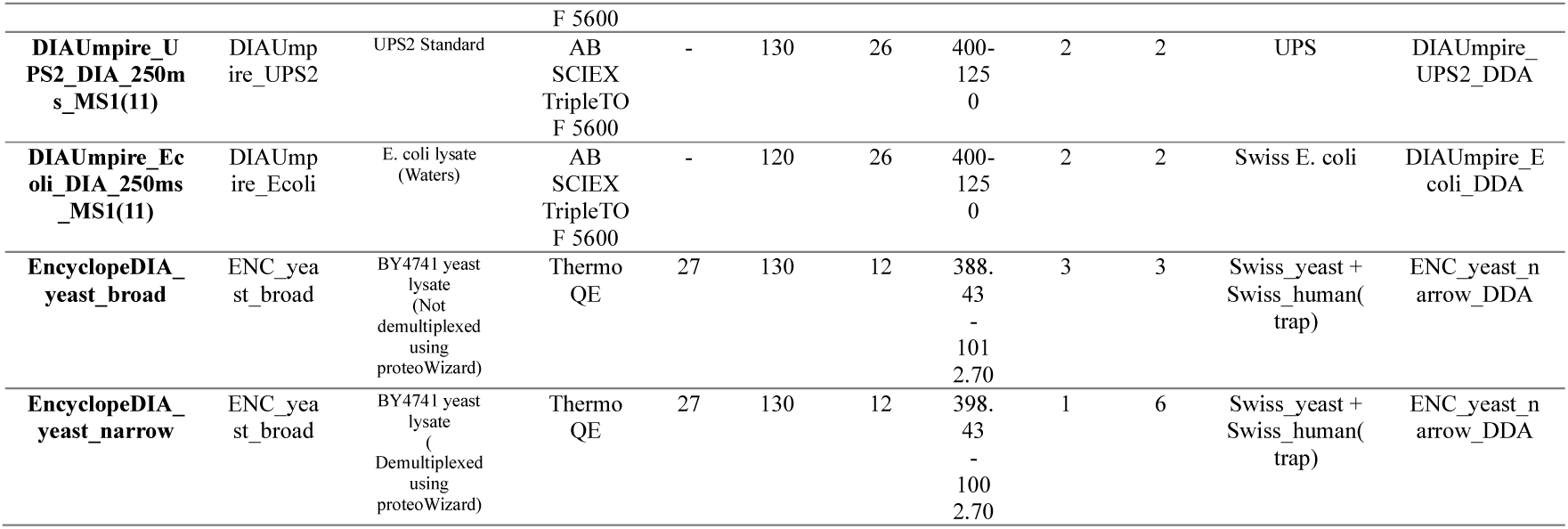
Dataset Details

**Supplementary Table 2.** Database Details

| Database | Description | #Proteins | File Size |
| --- | --- | --- | --- |
| **UPS** | Human | 50 | 16.0KB |
| **Swiss E. coli** | E. coli, Reviewed, Accessed at 2019-10-01 | 4518 | 1.84MB |
| **Swiss_human** | Human, Reviewed, Accessed at 2019-10-01 | 20431 | 13.0MB |
| **Swiss_Yeast** | Yeast, Reviewed, Accessed at 2019-10-01 | 20198 | 11.9MB |

## Footnotes

1 We achieve this by set max_round=1 (default settings: max_round=10). we have tried set max_round=0, but it spends more time since Open-pFind do not restrain the generation space of modification, which cause Open-pFind spent more time than max_round=1.

## References

1. Venable, J. D.; Dong, M. Q.; Wohlschlegel, J.; Dillin, A.; Yates, J. R., Automated approach for quantitative analysis of complex peptide mixtures from tandem mass spectra. Nat Methods 2004, 1, (1), 39–45.

2. Gillet, L. C.; Navarro, P.; Tate, S.; Rost, H.; Selevsek, N.; Reiter, L.; Bonner, R.; Aebersold, R., Targeted data extraction of the MS/MS spectra generated by data-independent acquisition: a new concept for consistent and accurate proteome analysis. Mol Cell Proteomics 2012, 11, (6), O111.016717.

3. Rost, H. L.; Rosenberger, G.; Navarro, P.; Gillet, L.; Miladinovic, S. M.; Schubert, O. T.; Wolski, W.; Collins, B. C.; Malmstrom, J.; Malmstrom, L.; Aebersold, R., OpenSWATH enables automated, targeted analysis of data-independent acquisition MS data. Nat Biotechnol 2014, 32, (3), 219–23.

4. MacLean, B.; Tomazela, D. M.; Shulman, N.; Chambers, M.; Finney, G. L.; Frewen, B.; Kern, R.; Tabb, D. L.; Liebler, D. C.; MacCoss, M. J., Skyline: an open source document editor for creating and analyzing targeted proteomics experiments. Bioinformatics 2010, 26, (7), 966–8.

5. Henderson, C. M.; Shulman, N. J.; MacLean, B.; MacCoss, M. J.; Hoofnagle, A. N., Skyline Performs as Well as Vendor Software in the Quantitative Analysis of Serum 25-Hydroxy Vitamin D and Vitamin D Binding Globulin. Clin Chem 2018, 64, (2), 408–410.

6. Pino, L. K.; Searle, B. C.; Bollinger, J. G.; Nunn, B.; MacLean, B.; MacCoss, M. J., The Skyline ecosystem: Informatics for quantitative mass spectrometry proteomics. Mass Spectrom Rev 2020, 39, (3), 229–244.

7. Demichev, V.; Messner, C. B.; Vernardis, S. I.; Lilley, K. S.; Ralser, M., DIA-NN: neural networks and interference correction enable deep proteome coverage in high throughput. Nat Methods 2020, 17, (1), 41–44.

8. Searle, B. C.; Pino, L. K.; Egertson, J. D.; Ting, Y. S.; Lawrence, R. T.; MacLean, B. X.; Villen, J.; MacCoss, M. J., Chromatogram libraries improve peptide detection and quantification by data independent acquisition mass spectrometry. Nat Commun 2018, 9, (1), 5128.

9. Ludwig, C.; Gillet, L.; Rosenberger, G.; Amon, S.; Collins, B. C.; Aebersold, R., Data-independent acquisition-based SWATH-MS for quantitative proteomics: a tutorial. Mol Syst Biol 2018, 14, (8), e8126.

10. Zhang, F.; Ge, W.; Ruan, G.; Cai, X.; Guo, T., Data-Independent Acquisition Mass Spectrometry-based Proteomics and Software Tools: A Glimpse in 2020. Proteomics 2020, e1900276.

11. Tsou, C. C.; Avtonomov, D.; Larsen, B.; Tucholska, M.; Choi, H.; Gingras, A. C.; Nesvizhskii, A. I., DIA-Umpire: comprehensive computational framework for data-independent acquisition proteomics. Nat Methods 2015, 12, (3), 258-64, 7 p following 264.

12. Zhou, X. X.; Zeng, W. F.; Chi, H.; Luo, C.; Liu, C.; Zhan, J.; He, S. M.; Zhang, Z., pDeep: Predicting MS/MS Spectra of Peptides with Deep Learning. Anal Chem 2017, 89, (23), 12690–12697.

13. Zeng, W. F.; Zhou, X. X.; Zhou, W. J.; Chi, H.; Zhan, J.; He, S. M., MS/MS Spectrum Prediction for Modified Peptides Using pDeep2 Trained by Transfer Learning. Anal Chem 2019, 91, (15), 9724–9731.

14. Searle, B. C.; Swearingen, K. E.; Barnes, C. A.; Schmidt, T.; Gessulat, S.; Kuster, B.; Wilhelm, M., Generating high quality libraries for DIA MS with empirically corrected peptide predictions. Nat Commun 2020, 11, (1), 1548.

15. Gessulat, S.; Schmidt, T.; Zolg, D. P.; Samaras, P.; Schnatbaum, K.; Zerweck, J.; Knaute, T.; Rechenberger, J.; Delanghe, B.; Huhmer, A.; Reimer, U.; Ehrlich, H. C.; Aiche, S.; Kuster, B.; Wilhelm, M., Prosit: proteome-wide prediction of peptide tandem mass spectra by deep learning. Nat Methods 2019, 16, (6), 509–518.

16. Sun, S.; Yang, F.; Yang, Q.; Zhang, H.; Wang, Y.; Bu, D.; Ma, B., MS-Simulator: predicting yion intensities for peptides with two charges based on the intensity ratio of neighboring ions. J Proteome Res 2012, 11, (9), 4509–16.

17. An, Z.; Zhai, L.; Ying, W.; Qian, X.; Gong, F.; Tan, M.; Fu, Y., PTMiner: Localization and Quality Control of Protein Modifications Detected in an Open Search and Its Application to Comprehensive Post-translational Modification Characterization in Human Proteome. Mol Cell Proteomics 2019, 18, (2), 391–405.

18. Searle, B. C.; Lawrence, R. T.; MacCoss, M. J.; Villen, J., Thesaurus: quantifying phosphopeptide positional isomers. Nat Methods 2019, 16, (8), 703–706.

19. Rosenberger, G.; Liu, Y.; Rost, H. L.; Ludwig, C.; Buil, A.; Bensimon, A.; Soste, M.; Spector, T. D.; Dermitzakis, E. T.; Collins, B. C.; Malmstrom, L.; Aebersold, R., Inference and quantification of peptidoforms in large sample cohorts by SWATH-MS. Nat Biotechnol 2017, 35, (8), 781–788.

20. Li, Y.; Zhong, C. Q.; Xu, X.; Cai, S.; Wu, X.; Zhang, Y.; Chen, J.; Shi, J.; Lin, S.; Han, J., Group-DIA: analyzing multiple data-independent acquisition mass spectrometry data files. Nat Methods 2015, 12, (12), 1105–6.

21. Tran, N. H.; Qiao, R.; Xin, L.; Chen, X.; Liu, C.; Zhang, X.; Shan, B.; Ghodsi, A.; Li, M., Deep learning enables de novo peptide sequencing from data-independent-acquisition mass spectrometry. Nat Methods 2019, 16, (1), 63–66.

22. Ting, Y. S.; Egertson, J. D.; Bollinger, J. G.; Searle, B. C.; Payne, S. H.; Noble, W. S.; MacCoss, M. J., PECAN: library-free peptide detection for data-independent acquisition tandem mass spectrometry data. Nat Methods 2017, 14, (9), 903–908.

23. Panchaud, A.; Scherl, A.; Shaffer, S. A.; von Haller, P. D.; Kulasekara, H. D.; Miller, S. I.; Goodlett, D. R., Precursor acquisition independent from ion count: how to dive deeper into the proteomics ocean. Anal Chem 2009, 81, (15), 6481–8.

24. Chi, H.; Liu, C.; Yang, H.; Zeng, W. F.; Wu, L.; Zhou, W. J.; Wang, R. M.; Niu, X. N.; Ding, Y. H.; Zhang, Y.; Wang, Z. W.; Chen, Z. L.; Sun, R. X.; Liu, T.; Tan, G. M.; Dong, M. Q.; Xu, P.; Zhang, P. H.; He, S. M., Comprehensive identification of peptides in tandem mass spectra using an efficient open search engine. Nat Biotechnol 2018.

25. Senko, M. W.; Beu, S. C.; McLaffertycor, F. W., Determination of monoisotopic masses and ion populations for large biomolecules from resolved isotopic distributions. Journal of the American Society for Mass Spectrometry 1995, 6, (4), 229–233.

26. Rockwood, A. L.; Haimi, P., Efficient calculation of accurate masses of isotopic peaks. J Am Soc Mass Spectrom 2006, 17, (3), 415–9.

27. Kall, L.; Canterbury, J. D.; Weston, J.; Noble, W. S.; MacCoss, M. J., Semi-supervised learning for peptide identification from shotgun proteomics datasets. Nat Methods 2007, 4, (11), 923–5.

28. Chen, T.; Guestrin, C. In Xgboost: A scalable tree boosting system, Proceedings of the 22nd acm sigkdd international conference on knowledge discovery and data mining, 2016; 2016; pp 785–794.

29. Friedman, J. H., Multivariate adaptive regression splines. The annals of statistics 1991, 1–67.

30. Michalski, A.; Damoc, E.; Hauschild, J. P.; Lange, O.; Wieghaus, A.; Makarov, A.; Nagaraj, N.; Cox, J.; Mann, M.; Horning, S., Mass spectrometry-based proteomics using Q Exactive, a high-performance benchtop quadrupole Orbitrap mass spectrometer. Mol Cell Proteomics 2011, 10, (9), M111.011015.

31. Chick, J. M.; Kolippakkam, D.; Nusinow, D. P.; Zhai, B.; Rad, R.; Huttlin, E. L.; Gygi, S. P., A mass-tolerant database search identifies a large proportion of unassigned spectra in shotgun proteomics as modified peptides. Nature biotechnology 2015, 33, (7), 743–749.

32. Sharma, K.; Schmitt, S.; Bergner, C. G.; Tyanova, S.; Kannaiyan, N.; Manrique-Hoyos, N.; Kongi, K.; Cantuti, L.; Hanisch, U.-K.; Philips, M.-A., Cell type–and brain region–resolved mouse brain proteome. Nature neuroscience 2015, 18, (12), 1819.

33. Amati, G., Two-Poisson model. In Encyclopedia of Database Systems, Liu, L.; ÖZsu, M. T., Eds. Springer US: Boston, MA, 2009; pp 3218–3219.

34. Teleman, J.; Rost, H. L.; Rosenberger, G.; Schmitt, U.; Malmstrom, L.; Malmstrom, J.; Levander, F., DIANA--algorithmic improvements for analysis of data-independent acquisition MS data. Bioinformatics 2015, 31, (4), 555–62.

35. Deutsch, E. W.; Mendoza, L.; Shteynberg, D.; Farrah, T.; Lam, H.; Tasman, N.; Sun, Z.; Nilsson, E.; Pratt, B.; Prazen, B.; Eng, J. K.; Martin, D. B.; Nesvizhskii, A. I.; Aebersold, R., A guided tour of the Trans-Proteomic Pipeline. Proteomics 2010, 10, (6), 1150–9.

36. Deutsch, E. W.; Mendoza, L.; Shteynberg, D.; Slagel, J.; Sun, Z.; Moritz, R. L., Trans-Proteomic Pipeline, a standardized data processing pipeline for large-scale reproducible proteomics informatics. Proteomics Clin Appl 2015, 9, (7-8), 745–54.

37. Liu, C.; Song, C. Q.; Yuan, Z. F.; Fu, Y.; Chi, H.; Wang, L. H.; Fan, S. B.; Zhang, K.; Zeng, W. F.; He, S. M.; Dong, M. Q.; Sun, R. X., pQuant improves quantitation by keeping out interfering signals and evaluating the accuracy of calculated ratios. Anal Chem 2014, 86, (11), 5286–94.

38. Ong, S. E.; Mann, M., A practical recipe for stable isotope labeling by amino acids in cell culture (SILAC). Nat Protoc 2006, 1, (6), 2650–60.

39. Li, W.; Chi, H.; Salovska, B.; Wu, C.; Sun, L.; Rosenberger, G.; Liu, Y., Assessing the Relationship Between Mass Window Width and Retention Time Scheduling on Protein Coverage for Data-Independent Acquisition. J Am Soc Mass Spectrom 2019, 30, (8), 1396–1405.

40. Eng, J. K.; Hoopmann, M. R.; Jahan, T. A.; Egertson, J. D.; Noble, W. S.; MacCoss, M. J., A deeper look into Comet--implementation and features. J Am Soc Mass Spectrom 2015, 26, (11), 1865–74.

41. Eng, J. K.; Jahan, T. A.; Hoopmann, M. R., Comet: an open-source MS/MS sequence database search tool. Proteomics 2013, 13, (1), 22–4.

